# Maize leaf functional responses to drought episode and rewatering

**DOI:** 10.1101/170258

**Authors:** He Song, Yibo Li, Li Zhou, Zhenzhu Xu, Guangsheng Zhou

## Abstract

Effects of crop growth and physiological activity to drought and irrigation regimes have been extensively studied; however, the responses of plant growth, morphological and photosynthetic behaviors to drought episodes and thereafter rewatering receive a less attention. This field experiment was carried out directly *in situ* at an agricultural ecosystem research station during 2015-2016, in a northeastern China, on the renowned northeastern maize production belt, where is being threatened by severe drought. A field automatic rain-shelter was used, and five irrigation regimes including control, four drought episodes, and rewatering treatments were established. The chlorophyll contents (SPAD values), light-saturated photosynthetic rate (*A*_sat_), and photosystem II actual quantum yield (Φ_PSII_), maximum quantum yield (*F*_v_′/*F*_m_′) decreased at lower leaf positions and with plant development. Episodic drought effects on plant growth, leaf morphological traits and photosynthetic processes at both vegetative and reproductive stages were severely remarked, particularly at late development stage and with longer drought duration. The recovery of leaf functional traits of the plants experienced historical-drought following re-irrigating was not fully restored to the level of the plants subjected to ample and normal water status; and the strength of recovery was proportional to the persistence of pre-drought episodes. The relationship of *A*_sat_ with SPAD depends on water status and plant development. A principal component analysis can well denote the change patterns in responses to water status treatments with plant development. The results may give an insight into how to understand the maize traits’ responses to drought episode and rewatering, and this also might assist the drought-stricken crops to cope with future climatic change.

## 1. Introduction

Climate change results in abnormal changes in precipitation patterns in terms of both its total amounts and episodic drought frequencies (Alley *et al.* 2003; Trenberth *et al.* 2014; IPCC 2014). Water shortage is a crucial constraint to crop growth, yield, physiological processes in many areas around world, including rain-fed and deficit-irrigation regions; meanwhile the abnormal occurrences of drought episodes usually fluctuates at various spatial-temporal scales (Boyer 1982; Battisti & Naylor 2009; IPCC 2014; Rurinda *et al.* 2014; Myers *et al.* 2017). Intensifying drought also may eliminate the expected benefits from some fewer favorite factors due to climate change such as elevated CO_2_ and enhanced anthropogenic nitrogen (N) deposit (Iversen & Norby 2014; Gray *et al.* 2016; Xu *et al.* 2016), and climatic warming may exacerbate drought disaster by further reducing soil moisture availability (Zeng *et al.* 2005; Lobell *et al.* 2011, 2014; Iversen & Norby 2014). As reported, due to potentially adverse climate change, since1950s to the present, agricultural drought-inducible disaster area also had an increasing trend in China—the drought-induced grain loss reached approximate 25-30 billion kg, accounting for 60% of total loss of natural disaster (Jiao *et al.* 2014, Zhou 2015). It has been notable that China’s agricultural drought becomes more serious mainly due to the adverse climate change and rapid social-economic development.

Maize is one of the most important three staple crops—maize, wheat, and rice, and the main resources of the feed, industrial raw materials (Campos *et al.* 2004; Long *et al.* 2006; Ribaut *et al.* 2009), recent years it ranked first place among the three staple crops (FAO 2017). In China, it also plays a critical role in food security and husbandry industry development among agricultural and even entire economic sectors at both regional and national levels (Meng *et al.* 2013; Ma & Ma 2017; PINC 2017). Drought is one of major limitations to maize production (Boyer 1982; Sharp *et al.* 2004; Xu *et al.* 2008; Lobell *et al.* 2014; Avramova *et al.* 2015), resulting in a yield reduction of 25-30%, even with no harvest in those years of extremely severe drought (Campos *et al.* 2004; Zhang *et al.* 2011). In USA major maize production zone, the drought sensitivity in maize production in recent two decades has been also reported to increase, despite cultivar improvements and the agronomic practices with higher sowing densities (Lobell *et al.* 2014). Climatic warming is projected to further exaggerate drought’s negative impact, leading to huge loss of maize production (Ribaut *et al.* 2009; Lobell *et al.* 2014). Drought stress leads to reductions in maize (*Zea mays* L.) and other crops’ yields mainly by (i) reducing plant growth and reproductive activities, (ii) reducing photosynthetic potentials and thereby radiation use efficiency (RUE), and (iii) reducing harvest index (HI) (Saini & Westgate 1999; Earl & Davis 2003; Barnabás *et al.* 2008; Xu *et al.* 2008). Contrastingly, if a maize cultivar root system and its ear growth are not completely limited, and leaf survival is enhanced despite water deficit, the cultivar may be recognized as high drought-tolerant one (Ribaut *et al.* 2009). Nevertheless, the intermittent drought imposition, and then following rewatering effects on crop plants grown in field still receive a relatively scant attention.

Based on cyclic drought experiment using *Catalpa bungei* species, the accumulative functional effects of progressive drought and subsequent re-watering on plant growth, leaf and root parameters has been found as a useful adaptive mechanism to drought successive drought and subsequent rewatering (Zheng *et al.* 2017). As recently reported by Abid *et al.* (2016), the adaptability to drought, and recovery rate and capacity was closely associated with wheat cultivars. The accumulation of effective metabolites such as sugars, and some amino acids like proline and leucine may exert an adaptive mechanism in response to the drought cyclic patterns (Meyer *et al.* 2014; Foster *et al.* 2015; Sun *et al.* 2016; Zheng *et al.* 2017). In plants of *Lupinus albus*, the new leaves can be produced more as quickly re-watered, although restoration of other metabolites (e.g. sugar content) was lagged (Pinheriro *et al.* 2004). Maize leaf length undergone one or several days of drought can restore completely following rewatering, but its growth rate could not reach the control level, suggesting that the growth resumption may be only a postponed event, no overcompensation occurrence (Acevedo *et al.* 1971). It is implied that the magnitude and rate of resumption might depend on pre-drought intensity and its duration (Hsiao 1973; Xu & Zhou 2007; Xu *et al.* 2009, 2010). Thus, the extent of compensation for the limitation of pre-drought by promoting plant growth as rewatering might determine the final plant biomass or crop yield, which may link to drought severity and its duration. Nevertheless, whether plant growth and physiological activities completely recovery following rewatering, what are the rate and degree of recovery, and the ability of the adaption to the drying-rewetting cycles might strongly depend on previous drought strength and persistent duration, species and genetic types, and drying-rewetting cycle patterns, which the underlying mechanism is elusive so far (Loewenstein & Pallardy 2002; Marron *et al.* 2003; Flexas *et al.* 2004; Yahdjian & Sala 2006; Xu *et al.* 2009, 2010; Sun *et al.* 2016; Zheng *et al.* 2017). Thus, responses of plant growth and leaf functional processes to drought history and subsequent rewatering remain to be clarified further, particularly *in situ* crop field.

As stated above, drought effects on plant/crop growth, photosynthesis, and other crucial eco-physiological process have been investigated extensively (e.g., Ne Smith & Ritchie 1992; Yordanov *et al.* 2002; Chaves *et al.* 2002; Harrison *et al.* 2014). However, as just recently stated by Abid *et al.* (2016), “studying plants’ capability to adapt and recover from drought stress is essential because of the ever-changing nature of drought events”. Herein, the objectives of this present study were to: (1) examine the effects of drought episode and rewatering on photosynthetic capacity and chlorophyll fluorescence; (2) compare the leaf functional responses to drought episode and rewatering at different leaf positions from bottom-most to upmost leaves, at various plant growth developments; (3) determine changing patterns in responses to drought episode and recovery after re-watering on photosynthetic capacity and chlorophyll fluorescence with the leaf developments; and (4) elucidate the index for drought adaptability and recovery ability following a pre-drought episode. Our hypotheses are expected: i) drought-episode-induced negative responses may depend on leaf ages/positions and leaf/plant development; ii) the amelioration of drought-induced negative responses by rewatering in field grown maize plants may mainly result from gas exchange behaviors relative to the chlorophyll fluorescence performances—photosystem II (PSII) photochemical processes; iii) the morphological and physiological functional traits may closely interacted, coordinately representing the adaptive responses to episodic drought and following re-wetting.

## 2. Materials and Methods

### 2.1. Site descriptions

The present two-year field experiment was carried out directly *in situ* at an agricultural ecosystem research station during 2015-2016 (41°49′N, 121°12′E, 27.4 m.a.s.l.), Jinzhou Ecology and Agricultural Meteorology Center, Jinzhou, Liaoning, a northeastern Chinese province on the renowned northeastern maize production belt (PINC 2017). This region is located in the northeast of the Eurasian areas, belongs to the warm temperate semi-humid monsoon climate, and atmospheric circulation mainly composed of westerlies and subtropical systems, with clear four seasons. The mean annual temperature is 7.8-9.0 °C, with the extreme maximum temperature of 41.8 °C and the extreme minimum temperature of −31.3 °C; annual frost-free period is 144-180 days; average annual rainfall is 540-640 mm, with 60% - 70% of rainfall concentrated in the summer. The soil is the typical brown soil, with a soil pH value of 6.3. The organic matter and total nitrogen content is 6.41-9.43 g kg^-1^ and 0.69 g kg^-1^, respectively. The staple crop in the region is maize (Han *et al.* 2007).

### 2.2. Experimental design

This study, a maize water-controlled field experiment, was conducted using a huge mobile rain-proof shelter during two growth seasons of 2015-2016. The two-year experimental design and its results were similar; thus, here the 2016-year results were mainly reported (for 2015-year experimental design and its results, see the Supporting Information File: Table S1 & S2, and Figures S1-S3). In the 2016-year experiment, the five irrigation treatments were designed: T_1_, T_2_, T_3_, T_4_, and T_5_ treatments, which denote Control, withholding water during jointing-tasseling, jointing-anthesis, tasseling-milking, and silking-milking, with 260, 188, 138, 136, and 161 mm irrigation amount in entire developmental stage, respectively (Table 1).

**Table 1.** Simulated rainfall regimes in the field experiment under a rain-shelter.

| Growth stage | Irrigation dates(Month/day) | T<sub>1</sub> | T<sub>2</sub> | T<sub>3</sub> | T<sub>4</sub> | T<sub>5</sub> |
| --- | --- | --- | --- | --- | --- | --- |
|  |  | Irrigating at every 7-day | Drought episode (27 days) | Drought episode (41 days) | Drought episode (41 days) | Drought episode (34 days) |
|  |  | Irrigation amount (mm) |  |  |  |  |
| V3~V13 | 5/24 | 8.7 | 8.7 | 8.7 | 8.7 | 8.7 |
|  | 5/30 | 0.8 | 0.8 | 0.8 | 0.8 | 0.8 |
|  | 6/8 | 10 | 10 | 10 | 10 | 10 |
|  | 6/15 | 10 | 10 | 10 | 10 | 10 |
|  | 6/22 | 10 | 10 | 10 | 10 | 10 |
|  | 6/29 | 24 | 24 | 24 | 24 | 24 |
| V13~VT | 7/6 | 24 | 0 | 0 | 24 | 24 |
|  | 7/13 | 24 | 0 | 0 | 24 | 24 |
|  | 7/20 | 25 | 0 | 0 |  | 25 |
| VT~R4 | 7/27 | 25 | 25 | 0 | 0 | 0 |
|  | 8/3 | 25 | 25 | 0 | 0 | 0 |
|  | 8/10 | 25 | 25 | 25 | 0 | 0 |
|  | 8/17 | 25 | 25 | 25 | 0 | 0 |
|  | 8/24 | 24 | 24 | 24 | 24 | 24 |
| Entire period | Total precipitation | 260.5 | 187.5 | 137.5 | 135.5 | 160.5 |

Notes: The dates measuring photosynthetic parameters are 1, July (V13, jointing); 14, July (V21); 25, July (VT, tasseling); 31, July (R1, Silking); 7, August (R2, blistering); 14, August (R3, milking); 29, August (R4, dough), respectively. Red parts indicate the drought episodes. Relative Soil water moisture (0 – 50 cm depth) decreased to a severe water deficit stress level of 30-40% at the ends of the drought episodes, meanwhile it maintained a normal level of 70-80% in the control plots or after rewatering. The maize growth stage nomenclature refers to Abendroth *et al.* (2009), http://lib.dr.iastate.edu/extension_pubs/26. Verified 21 July 2017.

There were three replicates in each treatment and 15 plots in total. Each plot is 5 m long and 3 m wide, surrounded by cement layer to avoid water permeation. The large mobile water-proof shelter is 4 m high, which is used for simulated precipitation to avoid the rainfall entrance. Maize cultivar used in this experiment was Danyu 405, which has been planted widely in this region. Seeds were sowed on 23 May, 2016. Controlled release fertilizer was used with 600 kg km^-2^.

### 2.3. Environmental variables and maize traits measurements

**Soil relative water content (SRWC) measurements** We used weighing method to measure the soil relative water content. Methods with soil auger were used to retrieve soil samples (0-50 cm), then put the samples to the aluminum specimen box, and weighed the samples to obtain the wet weight. Later, the samples were dried in an oven at 105 °C until a constant weight, and then the dried soil sample was weighed. There were three replicates in each treatment. The SRWC was calculated by the equation below:

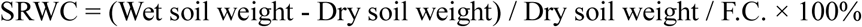

where F.C. is the soil water content measured 24 h after amply wetting the soil.

#### Chlorophyll content measurements

The relative chlorophyll contents (i.e., SPAD values) of maize leaves were measured by a SPAD-502 meter (Minolta Camera Co. Ltd., Japan). We chose 3-5 plants grown healthily for each treatment, and each leaf was measured three times on the leaf middle area to avoid the main vein, and then averaged.

#### Photosynthetic and chlorophyll fluorescence parameters

When measured the photosynthetic parameters, three-five plants grown healthily were chosen for each treatment, with a CIRAS-2 gas exchange system (PP Systems, Hertfordshire, UK). We measured the upper (i.e., the youngest and expanded leaves, 1–2 leaves from the top of the plants), middle (ear leaf or the leaves above or below ear leaf) and lower positions (relative orderly leaves) of each plant, respectively. Instrumentation system provided the red and blue built-in light source, and light intensity (Photosynthetically active radiation, PAR) was set to 1500 μmol m^-2^s^-1^. To ensure ample temperature and humidity conditions, the photosynthetic parameters of the maize leaves were measured at 9:00-11:00. The whole measuring process was used an open gas path, air relative humidity was controlled in 50% - 70%, CO_2_ concentration was controlled in 380 – 390 μmol mol^-1^, and leaf temperature was set up at around 27 °C. The parameters included light-saturated net photosynthetic rate (*A*_sat_), transpiration rate (*E*), stomatal conductance (*g_s_*); and the leaf water use efficiency (WUE) was calculated by the formula: WUE = *A*_sat_ / *E*.

The chlorophyll fluorescence parameters were measured using a chlorophyll fluorescence module (CFM) integrated with the CIRAS-2 gas exchange system at the same part of the same leaves measured simultaneously for the gas exchange parameters. First, the leaves were lighted at a light intensity of 1500 μmol m^-2^ s^-1^ after 15 min to measure the steady-state fluorescence (*F_s_*), and then gave a strong flash (5100 μmol m^-2^ s^-1^, with pulse time of 0.3 s) to measure the maximum fluorescence (*F*_m_′); later put the leaves under dark adaptation for 3 s, opened the far red after 5 s to measure the minimum light fluorescence (*F*_o_′). According to the expressions, we calculated chlorophyll fluorescence parameters: maximum quantum yield of PSII photochemistry (*F*_v_′*F*_m_′), quantum yield of PSII electron transport (Φ_PSII_), photochemical quenching (qP), and non-photochemical quenching (NPQ) (Genty *et al.* 1989; van Kooten & Snel 1990; Maxwell & Johnson 2000; Kramer *et al.* 2004):

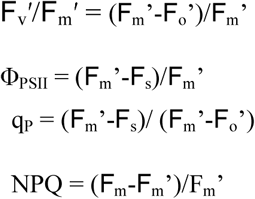

#### Plant height and leaf area measurements

Plant height and leaf area of maize were measured at different stages, and the measured dates were approximately consistent with those for the measurements of photosynthetic and fluorescence parameters. Maximum length and width were measured for ech leaf of a maize plant. A conventional formula of the total leaf area per plant was used (Francis *et al.* 1969):

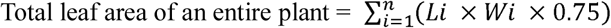

where *i* is the leaf order number of the measured plant, *n* is total number of the plant, *Li* is leaf maximum, and *Wi* leaf maximum width.

#### Leaf rolling index, leaf erection index, and leaf drooping angle determinations

The upper leaf actual width (at natural state, Ln), maximum width (at unfolding state, Lw), natural length (Lnl), maximum length (Lsl), basic angle of leaf (the angle between leaf and stem), drooping angle (under the naturally bending down of the leaf, the angle between the line from the pulvinus to the tip and the stem) were determined. Then, the leaf rolling index (LRI, %), leaf erection index (LEI, %), leaf bend degree (LBD, °) were calculated based on Xiang *et al.* (2012):

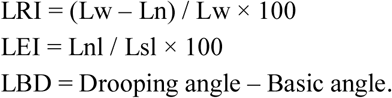

#### The plant biomass and grain yield

At the end of grain-filling, the tagged plants for measurements of these traits were retrieved, dried at 80°C to constant weight in a drying oven, and weighed to obtain the plant root, stem, leaf, and grain biomass.

### 2.4. Data statistics

The statistical analysis was used with SPSS 20.0 software (SPSS Inc., Chicago, IL). One-way ANOVA with Duncan’s multiple comparison was used to test the differences in the plant and leaf functional traits and morphological indicators between watering treatments at a 0.05 significance level. The differences of functional parameters among watering treatments, leaf positions and measurement dates were tested by the three-way ANOVA at the 0.05 significance level. The correlations between leaf functional and morphological traits were analyzed with Pearson method. The comprehensive patterns of the responses of leaf functional and morphological traits to episodic drought and rewatering were further analyzed with principal component analysis (PCA, Jolliffe 2002).

## 3. Results

### 3. 1. Photosynthetically physiological responses to drought episode and rewatering

#### 3.1.1 Responses in upper leaves

Photosynthetic physiological parameters were measured during the entire growing season: on 1, July (V13, jointing); 14, July (V21); 25, July (VT, tasselling); 31, July (R1, Silking); 7, August (R2, blistering); 14, August (R3, milking); 29, August (R4, dough), respectively (Table 1). As shown in Figure 1, the upper leaf net light-saturated photosynthetic rate (*A*_sat_) of control treatment (T_1_) increased with earlier plant development, reaching the maximum of 45.4 μmol m^-2^s^-1^ on 31 July (R1 stage), thereafter sharply decreasing until to a lowest point of 14.1 μmol m^-2^s^-1^ by 68.5% at later grain-filling stage.

**Figure 1.**
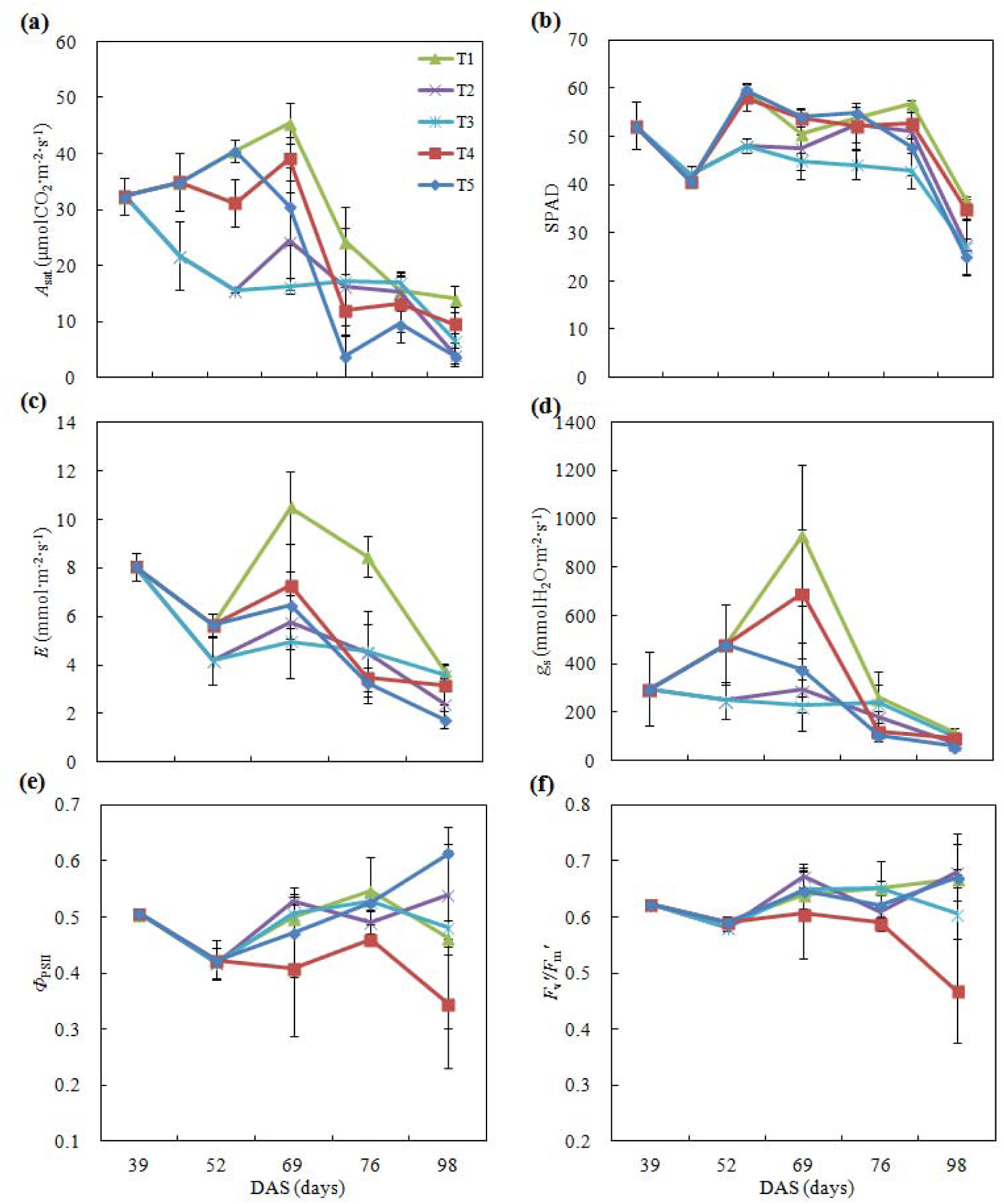
Effects of drought episode and rewatering on photosynthetic physiological processes in upper leaves. Notes: DAS, days after sowing; *A*_sat_, light-saturated photosynthetic rate; SPAD, relative chlorophyll contents; *g_s_*, stomatal conductance; *E*, transpiration rate; Φ_PSII_, PSII actual quantum yield; *F*_v_′/*F*_m_′, maximum quantum yield. T1, T2, T3, T4, and T5 denote Control, withholding water during jointing-tasseling, jointing-anthesis, tasseling-milking, and silking-milking, with 260, 188, 138, 136, and 161 mm irrigation amount in entire plant development, respectively.

For the T_2_ treatment (withholding water during jointing – tasseling, 27 days), the episodic drought led to a significant decline of *A*_sat_ with a low level of 21.7 μmol m^-2^s^-1^ on 14, July (V21 stage), a 40.3% drop; and further down to a lower level of 15.6 μmol m^-2^s^-1^ on 25, July (VT stage), a 65.6% drop. Upon re-irrigating, it rose up to 24.4 μmol m^-2^s^-1^ by 56.4% relative to the previous value, but not to reach the level of control treatment at the same stage—a maximum of 45.4 μmol m^-2^s^-1^ for the control plant. It indicated that a 27-day episodic drought from jointing to tasseling significantly inhibited photosynthetic capacity, just a part of recovery when following rewatering occurred, resulting in limitation to plant growth and development, and final grain yield loss.

Leaf *A*_sat_ of the T_3_ treatment (withholding water during jointing –anthesis, 41 days) always markedly decreased, even it remained a low and stable level as rewatering and during anthesis, thereafter declined rapidly after milking until to a low level of 6.6 μmol m^-2^s^-1^ at later grain-filling. The rewatering had a relatively-higher stimulation relative to the control only at milking, indicating the mild stimulation appeared (17.0 vs: 15.4 μmol m^-2^s^-1^, Figure 1a).

While leaf *A*_sat_ of the T_4_ treatment (withholding water during tasseling–milking, 41 days) had an increase at earlier withholding water (silking stage), then sharply declined to a low level of 12.0 μmol m^-2^s^-1^ at blistering, thereafter remaining a lower and stable level until the end of the grain-filling, although this time it seemed to lightly rise relative to T_3_ treatment, indicating that a stimulation occurred by the just nearly rewatering. Leaf *A*_sat_ of the T_5_ treatment (withholding water during silking–milking, 34 days), similar to T_4_ but less 7-day drought duration, declined even more sharply. It may indicate that later drought at tasseling may result in more sensitive effect on the gas exchange processes.

Other photosynthetically physiological parameters in the upper leaves showed similar change patterns (Figure 1b-f): For T_2_ and T_3_ plants, SPAD, *g*_s_, and *E* markedly deceased 4 weeks after withholding water. However, Φ_PSII_ and *F*_v_′/*F*_m_′ remained stable. Rewatering for T_2_ on 27, July, and for T_3_ on 10, August, resulted in stimulations in SPAD, *g_s_*, *E*, Φ_PSII_, and *F*_v_′/*F*_m_′; However, they still could not fully recover to the normal level. The drought stresses of T_4_ and T_5_ treatments also led to declines in the gas exchange rates, while the chlorophyll fluorescence parameters remained relatively stable except T_4_ treatment decrease during drought episode and thereafter rewatering.

#### 3.1.2 Responses in middle leaves

The changes in photosynthetic capacity of leaves at the plant middle position exhibited a similar change pattern in the upper leaves (Figure 2): As compared to the upper leaves, however, there were lower levels of *A*_sat_, SPAD, *g_s_*, and *E* during drought episodes, while rewatering following the pre-drying also did not lead to a complete restoration in the gas exchange parameters and SPAD values at later grain-filling stage. The chlorophyll fluorescence parameters still maintained a high level, implying that relatively mature middle leaves may have a drought-resistance in terms of PSII photochemical activities as compared with the younger upper leaves.

**Figure 2.**
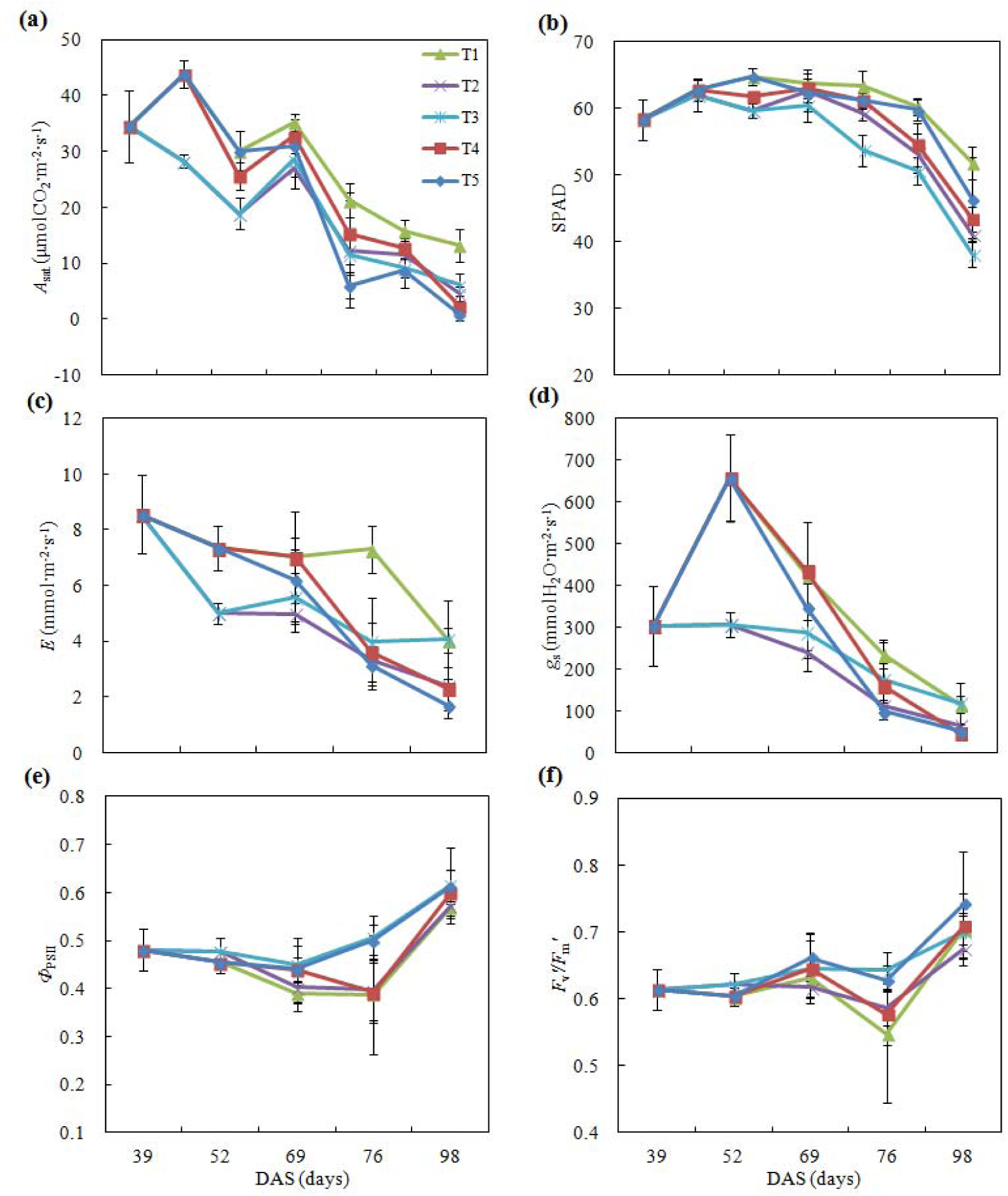
Effects of drought episode and rewatering on photosynthetic physiological processes in middle leaves. Notes: DAS, days after sowing; *A*_sat_, light-saturated photosynthetic rate; SPAD, relative chlorophyll contents; *g_s_*, stomatal conductance; *E*, transpiration rate; Φ_PSII_, PSII actual quantum yield; *F*_v_′/*F*_m_′, maximum quantum yield. T1, T2, T3, T4, and T5 denote Control, withholding water during jointing-tasseling, jointing-anthesis, tasseling-milking, and silking-milking, with 260, 188, 138, 136, and 161 mm irrigation amount in entire plant development, respectively

#### 3.1.3 Responses in bottom leaves

As measured in bottom leaves, there were greater levels in *A*_sat_ and SPAD under various water treatments at jointing stage (1, July), declining with plant developing (Figure 3). Plants at normal irrigation (control treatment) showed higher levels in the gas exchange parameters (*A*_sat_, *g_s_*, and *E*) and SPAD values, while lower levels were found during drought episodes, i.e., under T_3_, T_4_, and T_5_ treatments. Only part recovery was obtained upon rewatering at end of grain-filling stage in the plants experienced pre-drying history (T_4_ and T_5_ treatments). The chlorophyll fluorescence parameters such as Φ_PSI_ and *F*_v_′/*F*_m_′ were maintained at a higher level as the plants were subjected to withholding water treatments, particularly under T_3_ treatment (41-day withholding water during Jointing – anthesis). This again implicates an adaptive response to drought episode in terms of chlorophyll PSII photochemical activities.

**Figure 3.**
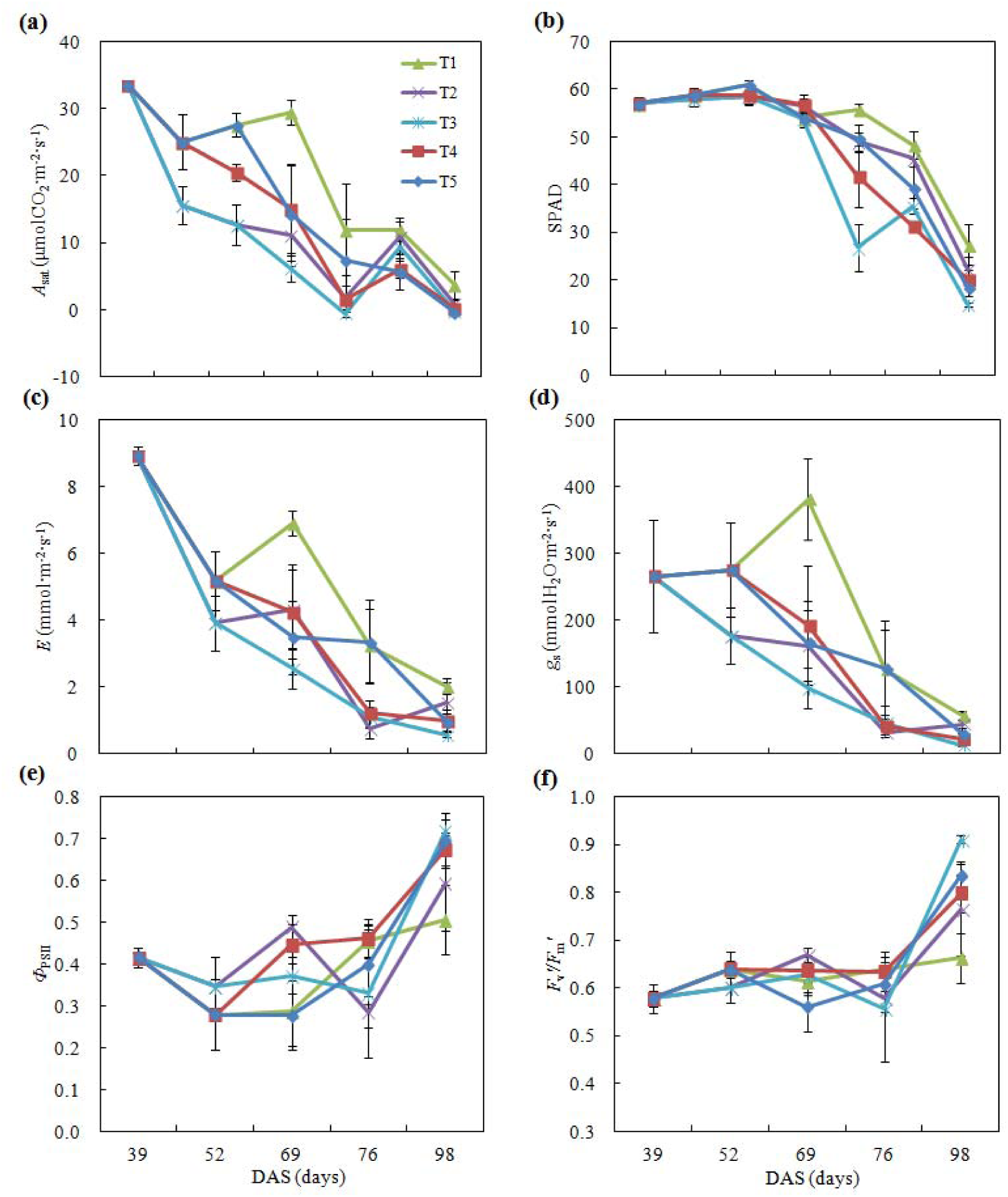
Effects of drought episode and rewatering on photosynthetic physiological processes in bottom leaves. Notes: DAS, days after sowing; *A*_sat_, light-saturated photosynthetic rate; SPAD, relative chlorophyll contents; *g_s_*, stomatal conductance; *E*, transpiration rate; Φ_PSII_, PSII actual quantum yield; *F*_v_′/*F*_m_′, maximum quantum yield. T1, T2, T3, T4, and T5 denote Control, withholding water during jointing-tasseling, jointing-anthesis, tasseling-milking, and silking-milking, with 260, 188, 138, 136, and 161 mm irrigation amount in entire plant development, respectively.

The three-way ANOVA showed that watering treatment, leaf position, and measured date produced significant effects on SPAD, *A*_sat_, *E* and WUE, individually (*P* < 0.001; Table 2). Only date alone and the interaction between leaf position and date had significant effects on *g_s_* (*P* < 0.05). The interactions between watering and date, and between leaf position and date significantly affected SPAD, *A*_sat_, and *E* (*P* < 0.001). Effect of leaf position and date alone, and their interaction on Φ_PSII_ were significant; while the significant effect on *F*_v_′/*F*_m_′ from date as single factor, and its interaction with leaf position appeared. The interaction of the three factors was not observed (*P* > 0.05).

**Table 2.** A significance list based on ANOVA for the effects and their interactions on the traits from watering, leaf position, and date

| | SPAD | $A_{\text{sat}}$ | $E$ | $g_s$ | WUE | $q_P$ | NPQ | $\Phi_{\text{PSII}}$ | $F_v'/F_m'$ |
| --- | --- | --- | --- | --- | --- | --- | --- | --- | --- |
| W | <0.001 | <0.001 | <0.001 | 0.988 | 0.007 | 0.114 | <0.001 | 0.215 | 0.946 |
| LP | <0.001 | <0.001 | <0.001 | 0.589 | <0.001 | <0.001 | 0.018 | 0.005 | 0.063 |
| Date | <0.001 | <0.001 | <0.001 | 0.009 | <0.001 | <0.001 | <0.001 | <0.001 | <0.001 |
| W×LP | 0.768 | 0.578 | 0.469 | 0.515 | 0.271 | 0.825 | 0.194 | 0.633 | 0.399 |
| W×D | <0.001 | 0.006 | <0.001 | 0.324 | <0.001 | 0.692 | <0.001 | 0.307 | 0.512 |
| LP×D | <0.001 | 0.002 | <0.001 | 0.014 | 0.454 | <0.001 | 0.178 | <0.001 | <0.001 |
| W×LP×D | 0.156ns | 0.963ns | 0.939ns | 0.946ns | 0.452ns | 0.692ns | 0.990ns | 0.850ns | 0.635ns |

Note: $P$ values are given. ns, no significant difference; W, LP, and D denote watering treatment, leaf position, and measurement data, respectively. ×, interaction sign; SPAD, relative chlorophyll content; $A_{\text{sat}}$ , light-saturated photosynthetic rate; $E$ , transpiration rate; $g_s$ , stomatal conductance; WUE, water use efficiency; $\Phi_{\text{PSII}}$ , PSII actual quantum yield; $F_v'/F_m'$ , maximum quantum yield; $q_P$ , photochemical quenching; NPQ, non-photochemical quenching.

The 2015-year experiment obtained the similar results on the responses of the relative chlorophyll content (SPAD values) and photosynthetic potentials mainly indicated by fluorescence parameters to the drought episode and the following rewatering regimes at the same experimental site (see Figures S1-S3).

### 3.2. Photosynthetically physiological responses in the leaves tagged at different developments

We measured the same leaves tagged to examine the photosynthetically physiological processes at different plant developments/leaf ages and the responses to episodic drought and rewatering—just staring from emerging of the leaves to becoming fully senesced. As showed in Figure 4, the leaf *A*_sat_ increased from the initial stage on first July, reaching a maximum on 25 July; thereafter, linearly declining with plant developing or leaf aging (Figure 4a). A gradual increase in SPAD values was observed initially with a maximum occurrence on 31 July, and a marked decline at end of grain-filling, i.e., 29 August (Figure 4b). Acute declines in *E* and *g_s_* were obtained after 31 July (Figure 4c,d); whereas the two chlorophyll fluorescence parameters (Φ_PSII_ and *F*_v_′/*F*_m_′) did not fluctuate greatly (Figure 4e,f).

**Figure 4.**
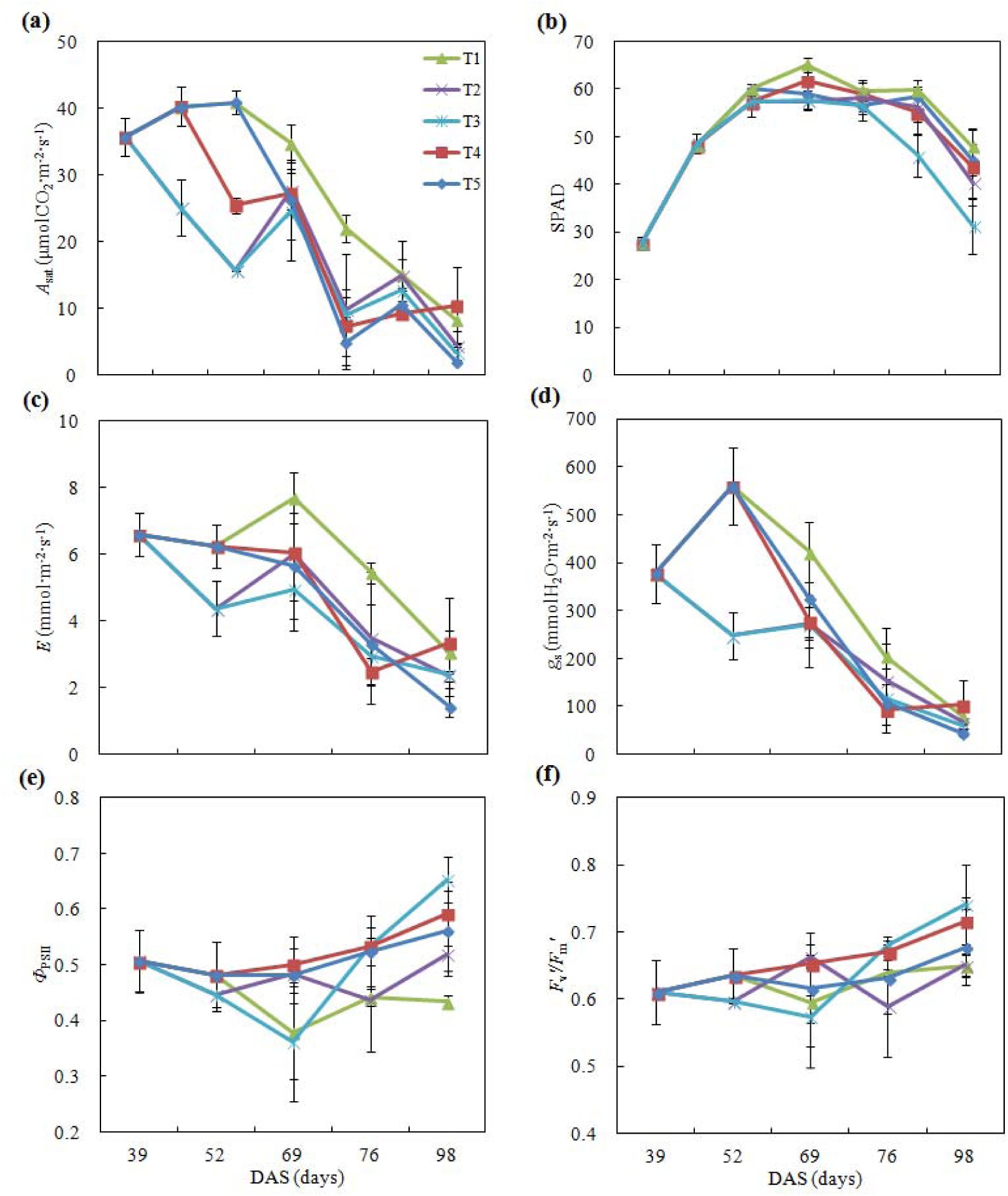
Photo synthetic physiological changes and the responses to drought episode and rewatering in the tagged leaves. Notes: DAS, days after sowing; *A*_sat_, light-saturated photosynthetic rate; SPAD, relative chlorophyll contents; *g_s_*, stomatal conductance; *E*, transpiration rate; Φ_PSII_, PSII actual quantum yield; *F*_v_′/*F*_m_′, maximum quantum yield. T1, T2, T3, T4, and T5 denote Control, withholding water during jointing-tasseling, jointing-anthesis, tasseling-milking, and silking-milking, with 260, 188, 138, 136, and 161 mm irrigation amount in entire plant development, respectively.

As compared to the normal water supply, drought episodes of T_3_, T_4_, and T_5_ treatments resulted in the declines in the tagged leaves’ gas exchange parameters—*A*_sat_, *g_s_*, and *E*, particularly with a longer drought persistence; while parts of stimulations by rewatering occurred after an episodic drought (Figure 4a,c,d). Decreases in SPAD values were often found as the drying episode occurred, particularly at the end of grain-filling (Figure 4b). We found that the two chlorophyll fluorescence parameters (Φ_PSII_ and *F*_v_′/*F*_m_′) had a slight response to either drought episodes or following rewatering, even with relative high levels upon the following rewatering (Figure 4e,f). This indicated that, in terms of chlorophyll fluorescence parameters, drought-tolerance may be enhanced with the leaf developing or its aging, this issue may need to be investigated further for different crops and their cultivars.

### 3.3. Responses of leaf morphological traits to drought and its relieving

Changes in maize plant canopy features (plant height, leaf area), leaf morphologic traits (leaf size, leaf rolling index (LRI), leaf erection index (LEI), leaf bend degree (LBD)) in responses to drought episodes and rewatering were shown in Table 3. The results at first measurement (1 July 2016) showed that no significant changes in plant height, plant individual leaf area, LRI, LEI, and LBD when the plants subjected to the normally watering treatments. LRIs of T_2_ and T_3_ on 14 July increased significantly due to their undergoing a 15-day drought episode duration. Plants of T_2_ and T_3_ reduced plant height and total leaf area, but increased LRI significantly on 25 July, indicating that the two drought treatments significantly affected plant and leaf growth, and morphological traits and canopy structure. LRI of T_3_ was greatest under drought on 31 July; and that of T_2_ just following rewatering rapidly decreased relative to the previous measurement; and the treatments of T_2_ and T_3_ had higher LBD. On 7 August, LRIs under drought conditions, i.e., T_3_, T_4_ and T_5_ treatments, were greater than that of the control, while LEI and LBD were not affected significantly. At milking stage (R3, On 14 August), LRI under T_4_ treatment was greater significantly, and that of T_2_ became lower close to control level due to rewatering. At the end of grain-filling stage (measured on 29 August), T_2_ plant leaf area was reduced significantly, whereas LRIs were not significant between the watering treatments. LEIs of T_4_ and T_5_ treatments were lower, indicating nearly rewatering did not trigger plant leaf erections with leaf aging.

**Table 3.** Effects of drought and rewatering on the plant growth and morphological indicators of maize (2016).

| Traits | Treatments | 1 Jul | 14 Jul | 25 Jul | 31 Jul | 7 Aug | 14 Aug | 29 Aug |
| --- | --- | --- | --- | --- | --- | --- | --- | --- |
| Plant height (m) | T<sub>1</sub> | 1.434±0.066a | 2.014±0.097a | **2.667±0.084a** | 2.717±0.119a | **2.734±0.134a** | 2.729±0.144a | 2.717±0.134a |
|  | T<sub>2</sub> | 1.336±0.057a | 1.897±0.051a | **2.233±0.093b** | 2.240±0.084a | **2.232±0.092b** | 2.428±0.224a | 2.214±0.108a |
|  | T<sub>3</sub> | 1.476±0.07a | 1.897±0.051a | **2.233±0.093b** | 2.327±0.132a | **2.436±0.148ab** | 2.407±0.157a | 2.333±0.199a |
|  | T<sub>4</sub> | 1.477±0.043a | 2.014±0.097a | **2.670±0.045a** | 2.710±0.147a | **2.745±0.104a** | 2.751±0.110a | 2.730±0.099a |
|  | T<sub>5</sub> | 1.355±0.009a | 2.014±0.097a | **2.667±0.084a** | 2.463±0.224a | **2.556±0.212ab** | 2.527±0.230a | 2.520±0.272a |
| Total leaf area (m<sup>2</sup> plant<sup>-1</sup>) | T<sub>1</sub> | 0.389±0.025a | 0.823±0.035a | **0.998±0.018a** | 1.049±0.021a | 1.037±0.044a | **0.974±0.017a** | **0.848±0.009a** |
|  | T<sub>2</sub> | 0.400±0.039a | 0.769±0.064a | **0.871±0.034b** | 0.890±0.065a | 0.930±0.047a | **0.809±0.038b** | **0.643±0.08b** |
|  | T<sub>3</sub> | 0.437±0.01a | 0.769±0.064a | **0.871±0.034b** | 0.953±0.097a | 0.871±0.054a | **0.686±0.042b** | **0.692±0.082ab** |
|  | T<sub>4</sub> | 0.423±0.038a | 0.823±0.035a | **0.988±0.023a** | 1.025±0.002a | 0.974±0.016a | **0.768±0.058b** | **0.781±0.012ab** |
|  | T<sub>5</sub> | 0.363±0.006a | 0.823±0.035a | **0.998±0.018a** | 0.877±0.078a | 0.879±0.067a | **0.837±0.07ab** | **0.738±0.043ab** |
| Leaf rolling index (LRI, %) | T<sub>1</sub> | 9.801±0.814a | **13.031±1.87b** | **7.993±2.824b** | **5.381±0.285c** | **9.712±1.127d** | **9.795±1.410c** | 12.907±0.682a |
|  | T<sub>2</sub> | 10.143±0.795a | **19.274±0.614a** | **17.626±1.356a** | **11.702±2.561b** | **12.126±2.358cd** | **10.453±0.910c** | 18.344±4.505a |
|  | T<sub>3</sub> | 11.116±0.288a | **19.274±0.614a** | **17.626±1.356a** | **23.194±1.653a** | **26.575±0.513a** | **17.24±1.212ab** | 18.692±1.192a |
|  | T<sub>4</sub> | 9.691±1.278a | **13.031±1.87b** | **9.594±1.546b** | **10.163±0.466b** | **16.867±0.658b** | **21.926±2.889a** | 18.93±3.082a |
|  | T<sub>5</sub> | 11.322±1.539a | **13.031±1.87b** | **7.993±2.824b** | **7.336±0.257bc** | **15.059±0.874bc** | **14.641±0.925bc** | 12.608±2.55a |
| Leaf erection index (LEI, %) | T<sub>1</sub> | 71.734±2.621a | 81.218±5.921a | 87.103±2.10a | 71.813±7.855a | 70.960±2.811a | 59.500±4.481a | **79.330±7.217a** |
|  | T<sub>2</sub> | 73.885±0.901a | 81.710±6.650a | 92.412±1.10a | 59.130±6.364a | 65.779±1.005a | 54.834±5.317a | **65.922±1.196a** |
|  | T<sub>3</sub> | 70.300±1.434a | 81.710±6.650a | 92.412±1.10a | 56.537±3.256a | 61.011±4.763a | 50.771±5.036a | **68.788±3.498a** |
|  | T<sub>4</sub> | 73.822±2.055a | 81.218±5.921a | 79.918±10.07a | 65.282±7.415a | 64.864±5.571a | 50.230±1.623a | **52.139±3.120b** |
|  | T<sub>5</sub> | 71.160±6.133a | 81.218±5.921a | 87.103±2.10a | 59.149±8.36a | 56.684±7.818a | 46.903±3.308a | **49.341±4.209b** |
| Leaf bend degree (LBD, °) | T<sub>1</sub> | 37.417±1.856a | 16.750±4.116a | 9.333±1.764a | **36.222±4.283c** | 53.567±6.19a | 81.417±9.419a | 59.983±20.037a |
|  | T<sub>2</sub> | 33.000±1.953a | 16.000±6.948a | 11.783±3.537a | **50.694±5.022ab** | 41.833±2.485a | 69.167±5.988a | 44.233±13.638a |
|  | T<sub>3</sub> | 39.667±5.516a | 16.000±6.948a | 11.783±3.537a | **55.833±4.757a** | 37.083±7.045a | 71.117±7.148a | 49.583±18.112a |
|  | T<sub>4</sub> | 33.917±1.609a | 16.750±4.116a | 13.833±2.167a | **31.722±3.692c** | 45.167±6.749a | 65.278±8.084a | 83.694±11.916a |
|  | T<sub>5</sub> | 35.750±3.527a | 16.750±4.116a | 9.333±1.764a | **37.944±1.811bc** | 50.733±2.709a | 60.833±6.194a | 55.361±4.835a |

Note: T<sub>1</sub>, T<sub>2</sub>, T<sub>3</sub>, T<sub>4</sub>, and T<sub>5</sub> denote Control, withholding water during jointing-tasseling, jointing-anthesis, tasseling-milking, and silking-milking, with 260, 188, 138, 136, and 161 mm irrigation amount in entire development, respectively. Different little-letters with mean data indicate significant differences between the five watering treatments (bold parts, n = 3-5).

### 4.4. Relationships between leaf morphological and functional traits in responses to drought and rewatering

Relationships between the morphological and functional traits of upper leaves were given in Table 4. The relationships among SPAD, *A*_sat_, *E*, *g_s_*, and WUE were significant (*P* < 0.05) except the correlations of WUE with *E* and *g_s_*. There were significant relationships between the two chlorophyll fluorescence parameters-Φ_PSII_ and *F*_v_′/ *F*_m_′ themselves; however, they did not correlate with SPAS values, and with gas exchange parameters (*P* > 0.05). Among plant morphological traits, there were significant and positive associations of plant height with total individual plant leaf area, LBD; but negative with LEI. LEI was significantly and negatively correlated with LBD. Between plant morphological and functional traits, there were significant and negative correlations of plant height with *A*_sat_, *g_s_*, WUE, and also those of LRI with SPAD, *A*_sat_, *E*, *g_s_* and WUE. Significant positive correlations of LEI with *A*_sat_ and *g_s_* were found, whereas significant negative correlations of LBD with SPAD, *A*_sat_, *g_s_* and WUE appeared, implicating the two leaf morphological indicators (i.e., LEI and LBD) may play an antagonistic role in the responses to drought episode and rewatering.

**Table 4.** Correlation coefficients between the functional and morphological traits of upper leaves

| | SPAD | $A_{\text{sat}}$ | $E$ | $g_s$ | WUE | $\Phi_{\text{PSII}}$ | $F_v'/F_m'$ | Plant height | Plant leaf area | LRI | LEI |
| --- | --- | --- | --- | --- | --- | --- | --- | --- | --- | --- | --- |
| $A_{\text{sat}}$ | **0.546\*\*** | | | | | | | | | | |
| $E$ | **0.498\*\*** | **0.813\*\*** | | | | | | | | | |
| $g_s$ | **0.427\*\*** | **0.845\*\*** | **0.816\*\*** | | | | | | | | |
| WUE | **0.304\*** | **0.479\*\*** | 0.042 | 0.186 |  |  |  |  |  |  |  |
| $\Phi_{\text{PSII}}$ | -0.152 | -0.184 | -0.039 | -0.143 | -0.206 | | | | | | |
| $F_v'/F_m'$ | -0.111 | -0.045 | 0.045 | -0.049 | -0.119 | **0.911\*\*** | | | | | |
| Plant height | -0.232 | **-0.391\*\*** | -0.168 | **-0.316\*\*** | **-0.344\*\*** | -0.036 | -0.02 |  |  |  |  |
| Plant leaf area | -0.021 | -0.211 | -0.035 | -0.222 | -0.238 | -0.015 | 0.021 | **0.854\*\*** |  |  |  |
| LRI | **-0.436\*\*** | **-0.506\*\*** | **-0.449\*\*** | **-0.422\*\*** | -0.204 | 0.014 | -0.039 | 0.18 | 0.094 |  |  |
| LEI | 0.145 | **0.328\*\*** | 0.217 | 0.264* | 0.227 | -0.166 | -0.025 | -0.298* | -0.185 | -0.211 |  |
| LBD | **-0.264\*\*** | **-0.383\*\*** | -0.232 | **-0.310\*** | **-0.279\*** | 0.119 | 0.007 | **0.389\*\*** | 0.139 | 0.151 | **-0.460\*\*** |

\* $P < 0.05$ , \*\* $P < 0.01$ represent significant differences. LRI, leaf rolling index; LEI, leaf erection index; LBD, leaf bend degree. Others see table 2.

To elucidate the critical linkage of the important functional traits such as relationship between the chlorophyll content and photosynthetic capacity in responses to watering treatment regimens at different growth stages, the correlations between SPAD and *A*_sat_ were analysed. The results showed that no significant relationship was observed from jointing to tasseling stages (Figure 5a), whereas significant-positive relationships appeared from silking to blistering (R^2^ = 0.12, *P* < 0.001), and from milking to denting (R^2^ = 0.41, *P* < 0.001; Figure 5b,c), indicating that their correlation becomes stronger with plant developmental process, particularly at later grain-filling. In addition, no significant relationship was found under normal irrigation condition (Figure 5d), meanwhile the significant relationship appeared under drought episode or rewatering treatments (Figure 5e,f), implying that their correlation can be enhanced due to the watering treatments.

**Figure 5.**
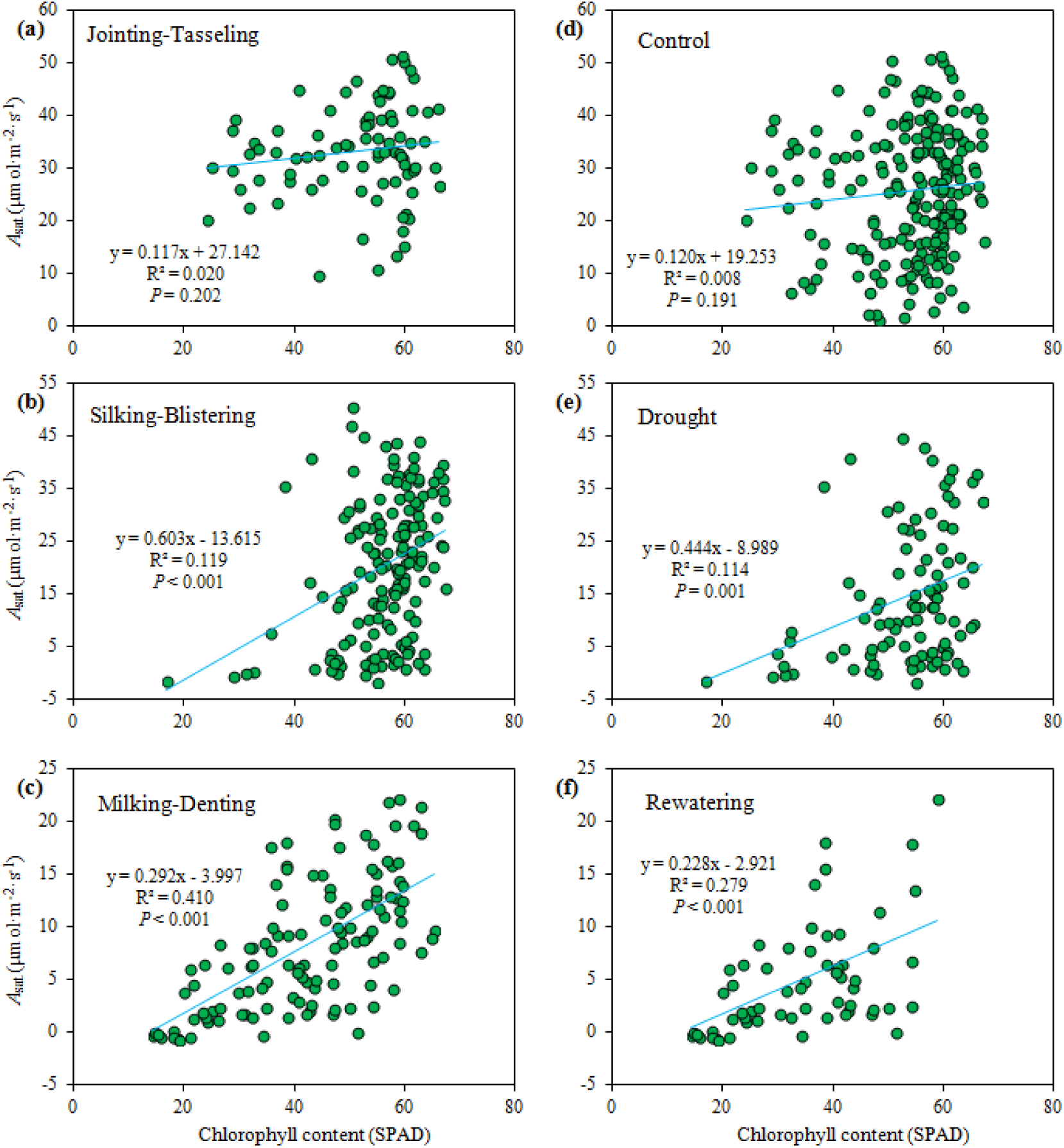
Effects of drought, rewatering, and plant developmental stages on relationships between light-saturated photosynthetic rate (*A*_sat_) and relative chlorophyll content (SPAD readings).

Based on the principal component analysis (PCA) on leaf functional and morphological traits at silking stage, the first two principal components (PCs) accounted for 60 % of the total variations of the maize traits (Figure 6). Gas exchange parameters closely positively correlated with principle component one (PC1), while the chlorophyll fluorescence parameters closely positively correlated with principle component two (PC2). There were negative correlations of PC2 with SPAD, plant height, and leaf area. The loadings were well typically distributed: the gas exchange parameters were located on the right side, the two chlorophyll fluorescence parameters upper part. SPAD was put alone on in quadrant III, while LRI was separated separately in quadrant II. Finally, the two plant growth traits ware located on quadrant IV, contrasting to LRI. Plots of the two factor scores demonstrated that the drought episode treatments (T_4_ and T_5_) were located on right-bottom parts, which is far from the loadings of gas exchange, chlorophyll fluorescence quantum yields. It implicates that the drought episode gave a severe impact on the photosynthetic capacity, and PCA may well represent the change patterns in plant and leaf growth, morphological and functional responses to the drought episode and rewatering regimes.

**Figure 6.**
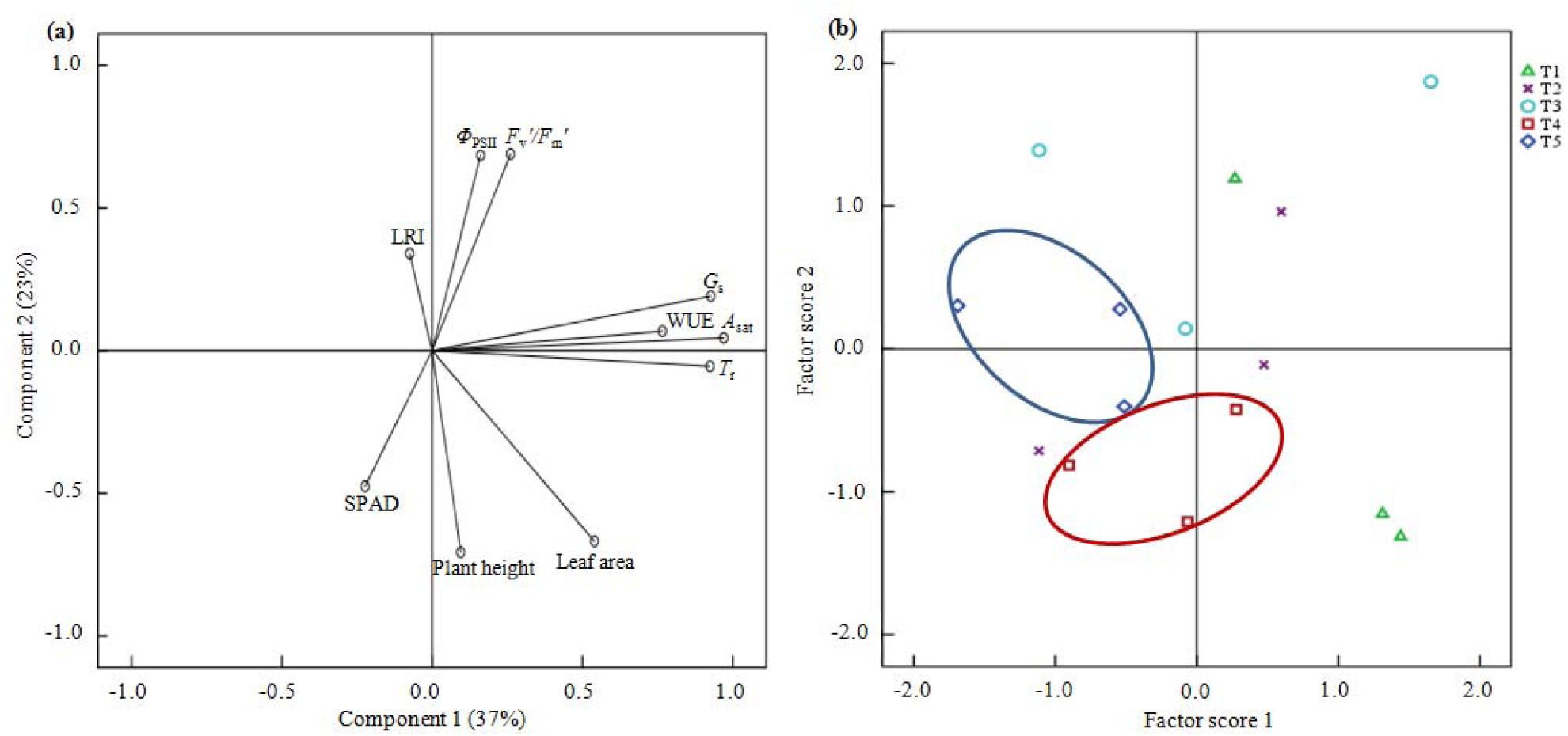
Principal component analysis on leaf functional and morphological traits and the effects of watering treatments at silking stage. Notes: LRI, leaf-rolling index; WUE, water use efficiency; red ellipse is for T_4_ treatment, while blue one is for T_5_ treatment. For others see Fig. 1 and table 3.

## Discussion

Maize, as one of most important staple crops, playing critical role in food security and husbandry industry development (Campos *et al.* 2004; FAO 2017), is being hampered severely by adverse climate change, such as abnormal precipitation alterations and heat wave events. Drought that is been intensifying by global warming is one of major limitations to maize growth and final grain yield (Boyer 1982; Sharp *et al.* 2004; Lobell *et al.* 2014; Jiao *et al.* 2014; Avramova *et al.* 2015; Myers *et al.* 2017). Although the effect on crop from drought as a single stress factor is studied extensively, understanding the crop functional processes in responses to drought episode and rewatering is still relatively scant, particularly across an entire plant development. The present results indicated that drought during jointing period produced severe effects on vegetative growth and photosynthetic capacity, while continuous drought during tasseling greatly-negatively impacted the reproductive growth and leaf photosynthetic capacity. Rewatering could alleviate adverse effects of pre-drought, but not stimulate both vegetative and reproductive growth and photosynthetic activities to recover fully to the levels of the control. Meanwhile, the leaf chlorophyll fluorescence parameters showed a relatively-stable and adaptive changes in response to episodic drying regimes. Moreover, the leaf functional traits mostly significantly negatively correlated with morphological indicators, and leaf rolling index (LRI) could be a sensitive indicator to assess the response of plants to drought and rewatering. Moreover, the association of light-saturated net photosynthetic rate (*A*_sat_) with chlorophyll content (SPAD value) can be as an effective proxy of leaf aging and the drought episodic durations. The present results may provide a newly profound insight into understanding the crops’ adaptive mechanism to drought cycle and rewatering regimes, and might be useful for the drought-resistant breeding practices and water-saving irrigation managements.

Nevertheless, the future models have well predicted that more extreme climate events will happen by the end of this century, making drought severer and heat waves more frequently (Luterbacher *et al.* 2004; Schär *et al.* 2004; IPCC 2014). The consequences of water availability loss have a marked influence on crop growth and productivity (Stuhlfauth *et al*. 1990; Ciais *et al.* 2005; Chaves *et al.* 2009), finally reducing the quantity and quality of grain yield (Marco & Tricoli.1993; Serraj & Sinclair. 2002; Barnabás et al. 2008; Lobell *et al.* 2014), and threatening food security regionally even globally (Ghannoum 2009; Myers *et al.* 2017). Among them, photosynthesis—the fundamental process for determination of crop growth and its development, even the final grain yield, is primarily affected by crucial climatic change factors such as drought (e.g., Boyer 1982; Chaves *et al.* 1991; Jefferies *et al.* 1994; Angelopoulos *et al*.1996; Farooq *et al.* 2009; Shen *et al.* 2015). The current results indicated that changes in the photosynthetically physiological parameters of the upper leaves are most significantly different between the different irrigation treatments (*P* < 0.05). Compared with control, under drought episode, the net light-saturated photosynthetic rate (*A*_sat_), chlorophyll content (SPAD value), stomatal conductance (*g_s_*), and transpiration rate (*E*), significantly decreased, while the actual quantum efficiency of PSII (Φ_PSII_), the effective quantum yield of PSII photochemistry (*F*_v_′/*F*_m_′) were no significantly declined. The previous experiments showed that the net photosynthesis rate decreases with *g_s_* dropping (Tenhunen 1987; Jarvis & Davies 1998; Miyashita *et al.* 2005). A decrease in *g_s_* will reduce water loss under drought stress, which also generates a drop in CO_2_ uptake (Frederick *et al.* 1989; Miyashita *et al.* 2005). When the plants of T_2_ treatment (pre-anthesis drought episode) was re-watered on 27 July, the photosynthetic parameters of leaves such as *A*_sat_, SPAD, *g_s_*, *E*, Φ_PSII_ and *F*_v_′/ *F*_m_′ mostly harmonically increased by the rewatering following the pre-drought episode; the similar responses to rewatering also occurred under T_3_, T_4_ and T_5_ treatments (Figures 1-3). As reported, precipitation events could generate a rapid response of the plant biological processes, which can trigger plants growth and development (e.g., Acevedo *et al.* 1971; Reynolds *et al.* 2004), but the values of photosynthetic parameters generally were lower than control level (Xu *et al.* 2009; Xu *et al.* 2010; Suralta *et al.* 2017). The results illustrated the continuous drought constrained photosynthetic capacity of the upper leaves, and the following rewatering is unable to totally recover to normal level (Figures 1 & 4). As reported previously, the degree and rate of recovery of rehydration might rely on the duration and severity of pre-drought (Xu & Zhou 2007; Chen *et al.* 2009; Xu *et al.* 2009, 2010; Abid *et al.* 2016).

The change patterns of middle leaves’ photosynthetic parameters were similar to those of the upper leaves (Figures 1-3): Under the normal treatment, SPAD, *A*_sat_, *g_s_*, *E*, remained at a higher level compared with drought episodes (T_3_, T_4_, and T_5_ treatments). Rewatering did not promote the photosynthetic capacities of T_4_, T_5_ at the grain-filling. In relation to the chlorophyll fluorescence parameters, which has been ones of the most practical and extensively used indicators to analysis plant eco-physiological processes (e.g., Maxwell & Johnson. 2000). Of them, *F*_v_′/*F*_m_′, as most useful one, often ranges between 0.80 and 0.83 without photoinhibition (Björkman & Demmig 1987). After suffering from severe water deficit stress, however, *F*_v_′/*F*_m_′ would decrease (Epron *et al.* 1992; Souza *et al.* 2004). However, our results showed both *F*_v_′/*F*_m_′ and Φ_PSII_ can maintain a high level even under severe drought. The results may explain that the middle leaves’ period of growth and development is longer, compared with the upper leaves. It may demonstrate a higher drought adaptability in the photochemical process and play a critical role in plant biomass accumulation and final grain yield (also see Allison & Watson 1966; Xu *et al.* 2008, 2011; Chen *et al.* 2016). For the bottom layer leaves, SPAD and *A*_sat_ were highest at the jointing stage (first measured on 1 July). Since then, they decreased with plant growing. The influence of drought on the bottom leaves was also consistent with the upper ones. SPAD and gas exchange parameters were also reduced markedly by the drought episode. At the end of grain-filling stage (24 August), the stimulating effect of rewatering was also lost partly. The two chlorophyll fluorescence parameters, however, remained higher under and after drought, especially at the end of grain-filling stage (e.g., withholding water from jointing to anthesis stages, total 41-day drought episode). It may again reflect the adaptive ability of maize elder leaves’ photochemical activity in responses to drought episodes and thereafter rewetting.

The results of the tagged leaves showed that along with the growth of the same leaf, the changing trends of the photosynthetic physiological parameters under each treatment were various (Figure 4): under control treatment, *A*_sat_ remained at a low level measured on 1 July, reaching a maximum on 25 July, thereafter linearly declining. SPAD reached a maximum on 31 July, thereafter significantly decreasing until grain-filling stage (29 August). Stomatal conductance and *E* decreased sharply after 31 July, whereas the changes in chlorophyll fluorescence parameters (Φ_PSII_, *F*_v_′/*F*_m_′) remained stable. The relative chlorophyll content and leaf gas exchange parameters of leaves in normally irrigated plants were at a high level, while those of the plants drought-stricken treatments (T_3_, T_4_ and T_5_ treatments) were at a low level, especially at the later grain-filling stage. It may highlight the leaf senescence process and its enhancement by drought stress (also see Xu *et al.* 2008; 2010). Rewatering can trigger gas exchange process; the chlorophyll fluorescence parameters (Φ_PSII_, *F*_v_′/*F*_m_′), however, were less affected by drought stress and rehydration (Lu & Zhang 1999; but Ghannoum *et al.* 2003; Gallé *et al.* 2007). Above all, the results again indicated that the drought-tolerance of the chlorophyll fluorescence parameters may increase with the plant/leaf growth and development when as measured in the same leaves (e.g., the tagged leaves). This issue remains debated, and needs the further research.

With the progresses of the maize plant and leaf growth, the photosynthetic performances of the upper, middle and lower leaves and the marked leaves of the maize may gradually decrease under moderate and severe drought stresses, and then recover partly after rewatering, which was consistent with previous research results (e.g., Flexas *et al.* 2009; Vaz *et al.* 2010; Xu *et al.* 2011). The current results also found that the photosynthetic performance of the middle leaves was higher than those of the both upper and lower leaves. The reasons may be due to the middle leaves being longer and greater than the upper one, and continuous drought may exacerbate further premature senescence of the bottom leaves (Iacono & Sommer 2000; Lu *et al.* 2001; He *et al.* 2002; Xu *et al.* 2011). Thus, the middle leave has stronger photosynthetic performances than both the upper and the lower ones, and it may contribute to most of carbon accumulation as plants exposed to drought episode and the following rewetting.

The sensitivity of crop yield to water deficits often markedly differs at different plant growth stages, which has been a classic study topic (Taylor *et al.* 1983; Fereres & Soriano 2007)—Generally, staple crops including maize are more sensitive to water deficit during seedling emergence, flowering, and early grain-filling than those during early plant growth and late grain-filling periods (e.g., Doorenbos & Kassam 1979). Under non-lethal water deficit during early vegetative growth, marked reductions in maize plant height and biomass were often found, and its crop phenology could delay, but this may not closely link to a lower yield potential (Damptey & Aspinall 1976; Abrecht & Carberry 1993). However, maize plant at jointing stage has exuberantly metabolic activities; under drought condition, the inhibition of plant height from jointing stage to tasseling stage was significantly greater than those at other stages (NeSmith & Ritchie 1992; Earl & Davis 2003). Therefore, the treatment with withholding water during the jointing period would mainly affect plant vegetative growth, leading to vegetative growth inhibition with short plant size, small leaves and internodes, and delays of tassel emergence, silk emergence, and onset of grain filling, finally a grain yield loss by 15-25% (Damptey & Aspinall 1976; NeSmith & Ritchie 1992). The current results indicated that the pre-anthesis drought significantly affected plant height and leaf size, even the canopy structure (and also see Earl & Davis 2003; Ne Smith & Ritchie 1992). Cakir (2004) found that the water content in the early growth stage of maize has great influence on plant height, consequently resulting in decline in canopy absorption of photosynthetically active radiation (PAR, Earl & Davis 2003). Thus, an early pre-anthesis water deficit might also not be neglected to obtain high yield even high in maize field, particularly with the severe and consecutive drought events (also see Ne Smith & Ritchie 1992). Our results manifested that the pre-anthesis drought markedly limited vegetative growth, lead to the declines in plant height and leaf area; and although the rehydration could alleviate the adverse effects from drought, it could not return fully to normal level at either early or later plant development stages. It highlights the drought constraints to maize plant growth during not only drought-persisting but drought-relieving periods. Additionally, leaf rolling index (LRI, Xiang *et al.* 2012), as a key parameter of plant morphological characteristics, obviously increased during drought episodes, but immediately decreased by rewatering. Therefore, it is suggested that the easy changes in LRI can be as a sensitive indicator for sensing drought and rewatering during almost entire plant growing season.

As stated above, drought stress seriously affected the photosynthesis of maize, inhibited the growth and development, and eventually leading to drops in plant biomass and yield (Irigoyen *et al*.1996; Bruce *et al.* 2002; Zhang *et al.* 2012). Maize grain yield is highly sensitive to drought during tasseling-silking period, mainly due to the marked decline in grain number (Otegui *et al.* 1995; Bolanos & Edmeades 1996; Barnabás *et al.* 2008). Our results also found that although the plants of T_4_ and T_5_ treatments were normally irrigated at jointing stage, the episodic drought at later tasseling stage, the ear growth was still significantly constrained relative to the control group. The current results of T_4_ and T_5_ treatments (pro-anthesis drought) showed no significant effect of drought at tasseling stage on the earlier plant growth. It again highlights that the later drought may exert a greatly adversely influence on the reproductive growth of maize plants.

The analysis on the correlation and principal component analysis reveal that there were positive correlations between the functional traits, except those with Φ_PSII_ and *F*_v_′/*F*_m_′; although a positive correlation themselves between Φ_PSII_ and *F*_v_′/*F*_m_′ was found. A significant-positive correlation between plant height and total leaf area, and a negative correlation between functional traits and morphological parameters occurred. SPAD is recognized an ideal non-destructive method for testing chlorophyll status in plant leaves, it has been widely used to investigate the environmental adaptability of crops (Manetas *et al.* 1998; Uddling *et al.* 2007; Steele *et al.* 2008; Ciganda *et al.* 2009; Pour-Aboughadareh *et al.* 2017). Interestingly, with the development of maize growth, the correlation of *A*_sat_ and SPAD was more positive and stronger, particularly under drought and rehydration conditions (Figure 5). Some results showed that the values of SPAD had a higher correlation with actual chlorophyll content under the conditions of lower actual chlorophyll content levels (Steele *et al.* 2008; Ciganda *et al.* 2009), depending on species and the chlorophyll distribution on leaf (Uddling *et al.* 2007). Therefore, probably maize plants grew at later growth stage and moisture condition became stressful, it may lead to reductions in both actual chlorophyll content and photosynthetic performance. Thus, the relationship between *A*_sat_ and SPAD might become more positive and closer (Figure 5). It is indicated that the relationships among the functional traits of leaves would alter with plant development and environmental changes, which may become closer and stronger with senescing at the later growth stage and especially under adverse environmental conditions.

In conclusion, from the current field experiment, episodic drought may exert markedly negative effects on photosynthetic potentials at either pre-anthesis or post-anthesis stages, particularly during early grain-filling; and the rapid recovery of leaf photosynthetic activities following re-irrigating occurred, but the recovery magnitude and rate might depend on the severity and persistence of the previous drought, and the leaf age and plant development. Generally, the leaf gas exchange, and leaf rolling index may demonstrate higher sensitivity to drought episode and thereafter rewatering than these chlorophyll content and its fluorescence parameters. The results would provide a profound insight into how to understand the crop functional traits’ responses to various drought stresses and precipitation or irrigation regimes. Finding the appropriate indicators for delayed leaf senescence while increased/stabilized photosynthetic potentials under changing water status may assist us in well dealing with climatic change to ensure crop production stability (Xu *et al.* 2010; Chen *et al.* 2013; Ben-Ari *et al.* 2016). Nevertheless, unambiguous understanding the underlying mechanisms of molecular and eco-physiological responses to various drought events, and helping the staple crops to cope with future climatic change to guarantee food security regionally and globally still remain a huge challenge, and that needs to be investigated further (Lobell *et al.* 2014; Gray *et al.* 2016; Myers *et al.* 2017).

## Acknowledgements

The study was funded by the National Natural Science Foundation of China (41330531), and China Special Fund for Meteorological Research in the Public Interest (GYHY201506019). The authors are grateful to Na Mi, Fu Cai, Kunqiao Shi, Yang Yang, Quanhui Ma for their work during the experiment.

## Availability of data and materials

The data sets supporting the results of this article are included within the article and its supporting information file.

## Authors’ contributions

ZX and GZ conceived and designed the study; HS, YL, LZ, ZX conducted the experiment and performed the data analysis; HS and ZX drafted the manuscript. All authors approved the final manuscript.

## References

Abendroth L, Elmore R, Hartzler RG, McGrath C, Mueller DS, Munkvold GP, Pope R, Rice ME, Robertson AE, Sawyer JE, Schaefer KJP, Tollefson JJ, Tylka GL. (2009) Corn Field Guide. Extension and Outreach Publications. Book 26. http://lib.dr.iastate.edu/extension_pubs/26. Verified 21 July 2017.

Abid M, Tian Z, Ata-Ul-Karim ST, Wang F, Liu Y, Zahoor R, et al. (2016). Adaptation to and recovery from drought stress at vegetative stages in wheat (*Triticum aestivum*) cultivars. Funct. Plant Biol. 43, 1159-1169.

Abrecht DG, Carberry PS. (1993). The influence of water deficit prior to tassel initiation on maize growth, development and yield. Field Crops Res. 31, 55-69.

Acevedo E, Hsiao TC, Henderson DW. (1971). Immediate and subsequent growth responses of maize leaves to changes in water statues. Plant Physiol. 48, 631–636.

Alley RB, Marotzke J, Nordhaus WD, Overpeck JT, Peteet DM, Pielke RA, et al. (2003). Abrupt climate change. Science, 299, 2005-2010.

Allison JCS, Watson DJ. (1966). The production and distribution of dry matter in maize after flowering. Ann. Bot. 30, 365–381.

Angelopoulos K, Dichio B, Xiloyannis C. (1996). Inhibition of photosynthesis in olive trees (*Olea europaea* L.) during water stress and rewatering. J. Exp. Bot. 47, 1093-1100.

Avramova V, AbdElgawad H, Zhang ZF, Fotschki B, Casadevall R, Vergauwen L, Knapen D, Taleisnik E, Guisez Y, Asard H, Beemster GTS. (2015). Drought induces distinct growth response, protection, and recovery mechanisms in the maize leaf growth zone. Plant Physiol. 169, 1382-1396.

Barnabás B, Jäger K, Fehér A. (2008). The effect of drought and heat stress on reproductive processes in cereals. Plant Cell Environ. 31, 11-38.

Battisti David S, Naylor Rosamond L. (2009). Seeds of doubt response. Science 324, 179-179.

Ben-Ari T, Adrian J, Klein T, Calanca P, Van der Velde M, Makowski D. (2016). Identifying indicators for extreme wheat and maize yield losses. Agr. Forest Meteorol. 220, 130-140.

Björkman O, Demmig B. (1987). Photon yield of O_2_ evolution and chlorophyll fluorescence characteristics at 77 K among vascular plants of diverse origins. Planta 170, 489-504.

Bolanos J, Edmeades GO. 1996. The importance of the anthesis-silking interval in breeding for drought tolerance in tropical maize. Field Crops Res. 48:65-80.

Boyer JS (1982) Plant productivity and environment. Science 218, 443-448.

Bruce WB, Edmeades GO, Barker TC. (2002). Molecular and physiological approaches to maize improvement for drought tolerance. J. Exp. Bot. 53, 13-25.

Çakir R. (2004). Effect of water stress at different development stages on vegetative and reproductive growth of corn. Field Crops Res. 89, 1-16.

Campos H, Cooper A, Habben JE, Edmeades GO, Schussler JR (2004) Improving drought tolerance in maize: a view from industry. Field Crop. Res. 90, 19-34.

Chaves MM, Flexas J, Pinheiro C. (2009). Photosynthesis under drought and salt stress: regulation mechanisms from whole plant to cell. Ann. Bot. 103, 551-560.

Chaves MM, Pereira JS, Maroco J, Rodrigues ML, Ricardo CPP, Osório ML, Pinheiro C. (2002). How plants cope with water stress in the field. Photosynthesis and growth. Ann Bot. 89, 907–916.

Chaves MM. (1991). Effects of water deficits on carbon assimilation. J. Exp. Bot. 42, 1-16.

Chen S, Lin G, Huang J, Jenerette GD. (2009). Dependence of carbon sequestration on the differential responses of ecosystem photosynthesis and respiration to rain pulses in a semiarid steppe. Glob. Change Biol. 15, 2450-2461.

Chen X, Chen F, Chen Y, Gao Q, Yang X, Yuan L, et al. (2013). Modern maize hybrids in Northeast China exhibit increased yield potential and resource use efficiency despite adverse climate change. Glob. Change Biol. 19, 923-936.

Chen Y, Wu D, Mu X, Xiao C, Chen F, Yuan L, Mi G. (2016). Vertical distribution of photosynthetic nitrogen use efficiency and its response to nitrogen in field-grown maize. Crop Sci. 56, 397–407.

Ciais P, Reichstein M, Viovy N, Granier A, Ogée J, Allard V, et al. (2005). Europe-wide reduction in primary productivity caused by the heat and drought in 2003. Nature 437, 529-533.

Ciganda V, Gitelson A, Schepers J. (2009). Non-destructive determination of maize leaf and canopy chlorophyll content. J. Plant Physiol. 166, 157-167.

Damptey HB, Aspinall D. (1976). Water deficit and inflorescence development in *Zea mays* L. Ann. Bot. 40, 23-35.

Doorenbos J, Kassam AH. (1979). Yield response to water. Irrigation and drainage paper 33:257.

Earl HJ, Davis RF. (2003). Effect of drought stress on leaf and whole canopy radiation use efficiency and yield of maize. Agron. J. 95, 688-696.

Epron D, Dreyer E, Breda N. (1992). Photosynthesis of oak trees [*Quercus petraea* (Matt.) Liebl.] during drought under field conditions: diurnal course of net CO_2_ assimilation and photochemical efficiency of photosystem II. Plant Cell Environ. 15, 809-820.

FAO. (2017). FAOSTAT. http://www.fao.org/faostat/en/#data. Verified 21 July 2017.

Farooq M, Wahid A, Kobayashi N, Fujita D, Basra SMA. (2009). Plant drought stress: effects, mechanisms and management. In Sustainable agriculture (pp. 153-188). Springer Netherlands.

Fereres E, Soriano MA. (2007). Deficit irrigation for reducing agricultural water use. J. Exp. Bot. 58, 147-159.

Flexas J, Barón M, Bota J, Ducruet JM, Gallé A, Galmés J, et al. (2009). Photosynthesis limitations during water stress acclimation and recovery in the drought-adapted vitis hybrid Richter-110 (V. *berlandieri* × *V. rupestris*). J. Exp. Bot. 60, 2361-2377.

Flexas J, Bota J, Loreta F, Cornic G, Sharkey TD. (2004). Diffusive and metabolic limitations to photosynthesis under drought and salinity in C_3_ plants. Plant Biol. 6, 1–11.

Francis CA, Rutger JN, Palmer AFE. (1969). A Rapid method for plant leaf area estimation in maize (*Zea mays* L.). Crop Sci. 9, 537-539.

Frederick JR, Alm DM, Hesketh JD. (1989). Leaf photosynthetic rates, stomatal resistances, and internal CO_2_ concentrations of soybean cultivars under drought stress. Photosynthetica 23, 575-584.

Gallé A, Haldimann P, Feller U. (2007). Photosynthetic performance and water relations in young pubescent oak (*Quercus pubescens*) trees during drought stress and recovery. New Phytol. 174, 799–810.

Genty B, Briantais J-M, Baker NR. (1989). The relationship between the quantum yield of photosynthetic electron transport and quenching of chlorophyll fluorescence. Biochimica et Biophysica Acta 990, 87-92.

Ghannoum O, Conroy JP, Driscoll SP, Paul MJ, Foyer CH, Lawlor DW. (2003). Nonstomatal limitations are responsible for drought induced photosynthetic inhibition in four C_4_ grasses. New Phytol. 159, 599–608.

Ghannoum O. (2009). C_4_ photosynthesis and water stress. Ann. Bot. 103, 635-644.

Gray SB, Dermody O, Klein SP, Locke AM, McGrath JM, et al. (2016). Intensifying drought eliminates the expected benefits of elevated carbon dioxide for soybean. Nature Plants 2, 16132. doi:10.1038/nplants.2016.132.

Han GX, Zhou GS, Xu ZZ, Yang Y, Liu JL, Shi KQ. (2007). Soil temperature and biotic factors drive the seasonal variation of soil respiration in a maize *(Zea mays* L.) agricultural ecosystem. Plant Soil 291, 15-26.

Harrison MT, Tardieu F, Dong Z, Messina CD, Hammer GL. (2014). Characterizing drought stress and trait influence on maize yield under current and future conditions. Glob. Change Biol. 20, 867-878.

He P, Osaki M, Takebe M, Shinano T. (2002). Changes of photosynthetic characteristics in relation to leaf senescence in two maize hybrids with different senescent appearance. Photosynthetica 40, 547–552.

Hsiao TC. (1973). Plant responses to water stress. Annu. Rev. Plant Physiol. 24, 519–570.

Iacono F, Sommer KJ (2000) Response of electron transport rate of water stress-affected grapevines: influence of leaf age. VITIS 39, 137–144.

IPCC. (2014). Climate Change 2014: Impacts, Adaptation, and Vulnerability. Part A: Global and Sectoral Aspects. Contribution of Working Group II to the Fifth Assessment Report of the Intergovernmental Panel on Climate Change, ed. CB Field, VR Barros, DJ Dokken, KJ Mach, MD Mastrandrea, et al. *Cambridge University Press*, *Cambridge, United Kingdom and New York*, *NY*, *USA*

Irigoyen JJ, Juan JP, Sanchez-Diaz MAN. (1996). Drought enhances chilling tolerance in a chilling-sensitive maize (Zea *mays)* variety. New Phytol. 134, 53-59.

Iversen C, Norby R. (2014). Terrestrial plant productivity and carbon allocation in a changing climate. In B. Freedman, Global Environmental Change (pp. 297-316). Springer Netherlands.

Jarvis AJ, Davies WJ. (1998). The coupled response of stomatal conductance to photosynthesis and transpiration. J. Exp. Bot., 49, 399-406.

Jefferies RA. (1994). Drought and chlorophyll fluorescence in field-grown potato (*Solanum tuberosum*). Physiol. Plant. 90, 93-97.

Jolliffe IT (2002) Principal component analysis. Springer, New York.

Jiao HY, Zhou GS, Chen ZL. (2014). Blue paper about agricultural issues on climate change. Assessment Report on Effect of Climate Change on China’ Agriculture (No.1). p1-6. Beijing: Social Science Literature Press.

Kramer DM, Johnson G, Kiirats O, Edwards GE. (2004). New fluorescence parameters for the determination of QA redox state and excitation energy fluxes. Photosynth. Res. 79, 209-218.

Lobell DB, Bänziger M, Magorokosho C, Vivek B. (2011). Nonlinear heat effects on African maize as evidenced by historical yield trials. Nat. Clim. Change 1, 42-45.

Lobell DB, Roberts MJ, Schlenker W, Braun N, Little BB, Rejesus RM, Hammer GL. (2014). Greater sensitivity to drought accompanies maize yield increase in the US Midwest. Science 344, 516-519.

Loewenstein NJ, Pallardy SG. (2002). Influence of a drying cycle on post-drought xylem sap abscisic acid and stomatal responses in young temperate deciduous angiosperms. New Phytol. 156, 351–361.

Long SP, Zhu X-G, Naidu SL, Ort DR. (2006). Can improvement in photosynthesis increase crop yields? Plant Cell Environ. 29, 315–330.

Lu C, Lu Q, Zhang J, Kuang T. (2001). Xanthophyll cycle, light energy dissipation and photosystem II down-regulation in senescent leaves of wheat plants grown in the field. Aust J. Plant Physiol. 28:1023–1030.

Lu C, Zhang J. (1999). Effects of water stress on photosystem II photochemistry and its thermostability in wheat plants. J. Exp. Bot. 50, 1199–1206.

Luterbacher J, Dietrich D, Xoplaki E, Grosjean M, Wanner H. (2004). European seasonal and annual temperature variability, trends, and extremes since 1500. Science 303, 1499-1503.

Manetas Y, Grammatikopoulos G, Kyparissis A. (1998). The use of the portable, non-destructive, SPAD-502 (Minolta) chlorophyll meter with leaves of varying trichome density and anthocyanin content. J. Plant Physiol. 153(3-4), 513-516.

Marco GD, Tricoli, D. (1993). Effect of water deficit on photosynthesis and electron transport in wheat grown in a natural environment. J. Plant Physiol. 142, 156–160.

Marron N, Dreyer E, Boudouresque E, Delay D, Petit J-M, Delmotte FM, Brignolas F. (2003). Impact of successive drought and re-watering cycles on growth and specific leaf area of two *Populus* × *canadensis* (Moench)clones,‘Dorskamp’ and ‘Luisa AvanzO’. Tree Physiol. 23, 1225–1235.

Ma X, Ma Y. (2017). The spatiotemporal variation analysis of virtual water for agriculture and livestock husbandry: A study for Jilin province in China. Sci. Total Environ. 586, 1150-1161.

Maxwell K, Johnson GN. (2000). Chlorophyll fluorescence—a practical guide. J. Exp. Bot. 51, 659-668.

Meng Q, Hou P, Wu L, Chen X, Cui Z, Zhang F. (2013). Understanding production potentials and yield gaps in intensive maize production in China. Field Crops Res. 143, 91-97.

Meyer E, Aspinwall MJ, Lowry DB, Palacio-Mejía JD, Logan TL, Fay PA, Juenger TE. 2014. Integrating transcriptional metabolomic, and physiological responses to drought stress and recovery in switchgrass *(Panicum virgatum* L.). BMC Genom. 15, 527–542.

Miyashita K, Tanakamaru S, Maitani T, Kimura K. (2005). Recovery responses of photosynthesis, transpiration, and stomatal conductance in kidney bean following drought stress. Environ. Exp. Bot. 53(2), 205-214.

Myers SS, Smith MR, Guth S, et al. (2017). Climate change and global food systems: potential impacts on food security and undernutrition. Annu. Rev. Public Health doi:10.1146/annurev-publhealth-031816-044356. ***In Press***.

Ne Smith DS, Ritchie JT. (1992). Short and long term responses of corn to a pre anthesis soil water deficit. Agron. J. 84, 107-113.

Otegui ME, Andrade FH, Suero EE. (1995). Growth, water use, and kernel abortion of maize subjected to drought at silking. Field Crops Res. 40, 87-94.

PINC. (2017). Crops Database. Planting Information Network of China. http://zzys.agri.gov.cn/nongqing.aspx, Verified, 21 July, 2017.

Pinheriro C, Passarinho JA, Ricardo CP. (2004). Effect of drought and rewatering on the metabolism of *Lupinus albus* organs. J Plant Physiol 161, 1203–1210

Pour-Aboughadareh A, Ahmadi J, Mehrabi AA, Etminan A, Moghaddam M, Siddique KH. (2017). Physiological responses to drought stress in wild relatives of wheat: implications for wheat improvement. Acta Physiol. Plant. 39, 106. doi:10.1007/s11738-017-2403-z. **In Press**.

Reynolds JF, Kemp PR, Ogle K, Fernández RJ. (2004). Modifying the ‘pulse–reserve’ paradigm for deserts of north America: precipitation pulses, soil water, and plant responses. Oecologia 141, 194-210.

Ribaut J-M, Betran J, Monneveux P, Setter T. (2009). Drought tolerance in maize. In: Bennetzen JL & Hake SC. (eds.), Handbook of Maize: Its Biology. p311-344. Springer Science + Business Media.

Rurinda J, Mapfumo P, van Wijk MT, Mtambanengwe F, Rufino MC, Chikowo R, Giller KE. (2014). Comparative assessment of maize, finger millet and sorghum for household food security in the face of increasing climatic risk. Eur. J. Agron. 55, 29-41.

Saini HS, Westgate ME. (1999). Reproductive development in grain crops during drought. Adv. Agron. 68, 59-96.

Schär C, Vidale PL, Lüthi D, Frei C, Häberli C, Liniger MA, et al. (2004). The role of increasing temperature variability in european summer heatwaves. Nature 427(6972), 332-6.

Serraj R, Sinclair TR. (2002). Osmolyte accumulation: can it really help increase crop yield under drought conditions? Plant Cell Environ. 25, 333-341.

Sharp RE, Poroyko V, Hejlek LG, Spollen WG, Springer GK, Bohnert HJ, Nguyen HT (2004) Root growth maintenance during water deficits: physiology to functional genomes. J. Exp. Bot. 55, 2343-2351.

Shen X, Dong Z, Chen Y. (2015). Drought and UV-B radiation effect on photosynthesis and antioxidant parameters in soybean and maize. Acta Physiol. Plant. 37, 1-8.

Souza RP, Machado EC, Silva JAB, Lagôa AMMA, Silveira JAG. (2004). Photosynthetic gas exchange, chlorophyll fluorescence and some associated metabolic changes in cowpea (*Vigna unguiculata*) during water stress and recovery. Environ. Exp. Bot. 51,45-56.

Steele MR, Gitelson AA, Rundquist DC. (2008). A comparison of two techniques for nondestructive measurement of chlorophyll content in grapevine leaves. Agron. J. 100, 779-782.

Stuhlfauth T, Scheuermann R, Fock HP. (1990). Light energy dissipation under water stress conditions contribution of reassimilation and evidence for additional processes. Plant Physiol. 92, 1053-61.

Sun C, Gao X, Chen X, Fu J, Zhang Y. (2016). Metabolic and growth responses of maize to successive drought and re-watering cycles. Agr. Water Manag. 172, 62-73.

Suralta RR, Kano-Nakata M, Niones JM, Inukai Y, Kameoka E, Tran TT, et al. (2017). Root plasticity for maintenance of productivity under abiotic stressed soil environments in rice: Progress and prospects. Field Crops Res. doi:10.1016/j.fcr.2016.06.023. ***In Press***.

Taylor HM, Jordan WR, Sinclair TR. (1983). Limitations to efficient water use in production. Madison, WIASA, CSSA, SSSA.

Tenhunen JD, Pearcy RW, Lange and OL. (1987). Diurnal variations in leaf conductance and gas exchange in natural environments. In: Zeiger, Farquhar GD & Cowan IR (eds), Stomatal Function, p323. Stanford University Press, Stanford, California.

Trenberth KE, Dai A, Van Der Schrier G, Jones PD, Barichivich J, Briffa KR, Sheffield J. (2014). Global warming and changes in drought. Nat. Clim. Change 4, 17-22.

Uddling J, Gelang-Alfredsson J, Piikki K, Pleijel H. (2007). Evaluating the relationship between leaf chlorophyll concentration and SPAD-502 chlorophyll meter readings. Photosynth. Res. 91, 37-46.

van Kooten O, Snel JFH. (1990). The use of chlorophyll fluorescence nomenclature in plant stress physiology. Photosynth. Res. 25, 147-150.

Vaz M, Pereira JS, Gazarini LC, David TS, David JS, Rodrigues A, et al. (2010). Drought-induced photosynthetic inhibition and autumn recovery in two Mediterranean oak species (*Quercus ilex* and *Quercus suber*). Tree Physiol. 30, 946-956.

Xiang JJ, Zhang GH, Qian Q, Xue HW. (2012). Semi-rolled leaf1 encodes a putative glycosylphosphatidylinositol-anchored protein and modulates rice leaf rolling by regulating the formation of bulliform cells. Plant Physiol. 159, 1488-1500.

Xu ZZ, Zhou GS. (2007). Photosynthetic recovery of a perennial grass *Leymus chinensis* after different periods of soil drought. Plant Prod. Sci. 10, 277-285.

Xu ZZ, Zhou GS, Han GX, Li YJ. (2011). Photosynthetic potential and its association with lipid peroxidation in response to high temperature at different leaf ages in maize. J. Plant Growth Regul. 30, 41-50.

Xu ZZ, Zhou GS, Shimizu H. (2010). Plant responses to drought and rewatering. Plant Signal. Behav. 5, 649-654.

Xu ZZ, Zhou GS, Shimizu H. (2009). Are plant growth and photosynthesis limited by pre-drought following rewatering in grass? J. Exp. Bot. 60, 3737–3749.

Xu ZZ, Zhou GS, Wang YL, Han GX, Li YJ. (2008). Changes in chlorophyll fluorescence in maize plants with imposed rapid dehydration at different leaf ages. J. Plant Growth Regul. 27, 83-92.

Xu ZZ, Zhou GS. (2007). Photosynthetic recovery of a perennial grass after different periods of soil drought. Plant Production Sci. 10, 277-285.

Yahdjian L, Sala OE. (2006). Vegetation structure constrains primary production response to water availability in the Patagonian steppe. Ecology 87, 952–962.

Yordanov I, Velikova V, Tsonev T. (2000). Plant responses to drought, acclimation, and stress tolerance. Photosynthetica 38, 171-186.

Zhang L, Gao M, Hu J, Zhang X, Wang K, Ashraf M. (2012). Modulation role of abscisic acid (ABA) on growth, water relations and glycinebetaine metabolism in two maize (*Zea mays* L.) cultivars under drought stress. Int. J. Mol. Sci. 13, 3189-3202.

Zhang RH, Xue JQ, Pu J, Zhao B, Zhang XH, Zheng YJ, Bu LD. (2011). Effects of drought stress on plant growth and photosynthetic characteristic of maize seedlings. Acta Agron. Sin. 37, 521-528.

Zheng H, Zhang X, Ma W, Song J, Rahman SU, Wang J, Zhang Y. (2017). Morphological and physiological responses to cyclic drought in two contrasting genotypes of *Catalpa bungei*. Environ. Exp. Bot. 138, 77–87.

Zhou GS. (2014). Research perspectives on effect of climate change on China’ Agricultural production. Meteorol. Environ. Sci. 38, 80-94.

